# Impacts of Recurrent Hitchhiking on Divergence and Demographic Inference in *Drosophila*

**DOI:** 10.1101/187633

**Authors:** Jeremy D. Lange, John E. Pool

## Abstract

In species with large population sizes such as *Drosophila*, natural selection may have substantial effects on genetic diversity and divergence. However, the implications of this widespread non-neutrality for standard population genetic assumptions and practices remain poorly resolved. Here, we assess the consequences of recurrent hitchhiking (RHH), in which selective sweeps occur at a given rate randomly across the genome. We use forward simulations to examine two published RHH models for *D. melanogaster*, reflecting relatively common/weak and rare/strong selection. We find that unlike the rare/strong RHH model, the common/weak model entails a slight degree of Hill-Robertson interference in high recombination regions. We also find that the common/weak RHH model is more consistent with our genome-wide estimate of the proportion of substitutions fixed by natural selection between *D. melanogaster* and *D. simulans* (19%). Finally, we examine how these models of RHH might bias demographic inference. We find that these RHH scenarios can bias demographic parameter estimation, but such biases are weaker for parameters relating recently-diverged populations, and for the common/weak RHH model in general. Thus, even for species with important genome-wide impacts of selective sweeps, neutralist demographic inference can have some utility in understanding the histories of recently-diverged populations.

## Introduction

The advancement of DNA sequencing technology, along with computational capacity and methodology, continues to revolutionize the field of population genetics. Harnessing the power of whole genome datasets, researchers have begun to explore a wider variety of evolutionary models. One such model that has received considerable attention recently is a model of recurrent hitchhiking, where genetic diversity at neutral regions is reduced due to repeated selective sweeps at nearby loci. This reduction in diversity has been explored theoretically (Kaplan *et al*. 1989; Stephan *et al*. 1992; Wiehe and Stephan 1993) showing that the expected reduction in diversity can be approximated as a function of RHH model parameters: *λ = 2N_e_s* and *λ*, where *N_e_* is the effective population size, s is the selection coefficient, and *λ* is the rate of positively selected substitutions. Subsequent studies have examined such RHH models using forward simulation, focusing attention on how Hill-Robertson interference (HRI; Hill and Robertson 1966) between linked beneficial mutations on different haplotypes reduces the probability of fixation (Gerrish and Lenski 1998, Chevin et al. 2008).

The impact of natural selection on genomic diversity may be particularly significant for species with very large population sizes, such as *Drosophila melanogaster* (*e.g*. Sella *et al*. 2009; Langley *et al*. 2012). In abundant taxa, the population adaptive mutation rate is elevated and the weak influence of genetic drift may allow natural selection to favor alleles with modest selection coefficients. By estimating RHH parameters, Jensen *et al*. (2008) suggested that selective sweeps may reduce genomic diversity in *D. melanogaster* to half of neutral levels. While this study implicated a model of relatively strong and infrequent sweeps, the study of Andolfatto (2007) instead favored a model of substantially weaker but more frequent adaptive substitutions. Though both of these studies utilized the same genomic data (synonymous polymorphism data at 137 X-linked loci in *D. melanogaster*) to infer selection strength and adaptive mutation rate, the methods of the two studies led to distinct conclusions. The Jensen study utilized an Approximate Bayesian Computation method to jointly infer adaptive mutation rate and selection strength. The Andolfatto study, however, used a maximum likelihood approach to estimate the product of the adaptive mutation rate and selection strength, followed by a McDonald-Kreitman approach to separate the two (McDonald and Kreitman 1991). While these models should imply strongly different proportions of substitutions driven by positive selection, their alignment with estimates of this quantity from genome-wide *Drosophila* data is unclear. And likewise, the predictions of each model for the role of HRI and for the fixation of neutral variants have not been investigated. We therefore sought to clarify the relationship between published *Drosophila* RHH models and adaptive divergence.

If linkage to natural selection substantially impacts *Drosophila* genetic diversity at neutral sites, the accuracy of demographic inference methods that assume neutrality is not assured. Most sites in the fly genome experience direct functional constraint (Halligan and Keightley 2006), which may lead to an excess of rare alleles from deleterious polymorphisms. Many sites that are not under as much direct selection pressure, such as synonymous sites and middles of short introns, are by definition very close to nonsynonymous sites and other functional sites that may experience natural selection. Selective sweeps could skew the genome-wide allele frequency spectrum, in particular by generating a skew toward rare alleles (Braverman *et al*. 1995) that may resemble the predictions of recent population growth. In line with these concerns, Schrider *et al*. (2016) found that the presence of positive selection can bias demographic parameter estimates for a single population’s history, and can lead to misidentification of demographic models. However, much interest centers on the inference of demographic parameters between recently-diverged populations, and it remains unclear whether Drosophila-like RHH on shorter time-scales is sufficient to bias parameters concerning population divergence times, population-specific size changes, and migration rates. We therefore use RHH simulations to investigate the impact of RHH on estimation of these parameters.

## Materials and Methods

### McDonald-Kreitman analysis

To estimate the proportion of substitutions in the *D. melanogaster* genome fixed by natural selection, we applied a genome-wide asymptotic McDonald-Kreitman analysis (Messer and Petrov 2013, McDonald and Kreitman 1991) using the web tool from Haller and Messer 2017. Here, we surveyed 197 genomes from a Zambian population of *D. melanogaster* (Lack *et al*. 2015), which is believed to be within the ancestral range of the species (Pool *et al*. 2012). These genomes are masked for identity by descent, apparent heterozygosity, and recent cosmopolitan admixture. In our analysis, we require any given site to be called in at least 50% of the genomes. We also applied a more conventional McDonald-Kreitman analysis, in which we required the minor allele to be segregating above 10% frequency in our sample. Applying this filter to both putatively neutral and selected site classes should reduce bias from deleterious polymorphisms. To estimate the number of substitutions, we used a *Drosophila simulans* genome aligned to the *D. melanogaster* genome (Stanley and Kulathinal 2016).

In this analysis, we estimated the proportion of substitutions driven to fixation by natural selection as 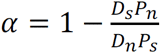 (Smith and Eyre-Walker 2002). Here, *D_s_* is the number of synonymous substitutions, *P_n_* is the number of nonsynonymous polymorphic sites, *D_n_* is the number of nonsynonymous substitutions, and *P_s_* is the number of synonymous polymorphic sites. *α* was calculated for nine site classes (nonsynonymous, two-fold synonymous, three-fold synonymous, 5’ untranslated regions, 3’ untranslated regions, intron, intergenic, and RNA-coding) and individually for each major chromosome arm (2L, 2R, 3L, 3R, and X). Four-fold synonymous sites were evaluated as proxies for neutral evolution. Site classes were taken from flybase.org for release 5.56 of the *D. melanogaster* genome.

### Simulations

In this study, we are interested in the effects of recurrent hitchhiking on demographic inference. To examine this, we ran forward simulations using SLIM version 2.5 (Haller & Messer 2016) to model recurrent hitchhiking. Because full-forward simulations are memory intensive and slow when simulating large populations, it is necessary to rescale simulation simulation parameters. We started by running test simulations to get an idea of the largest population size that we could simulate in a reasonable amount of time. We concluded that diploid populations of 50,000 individuals were a sensible target. This results in 50X rescaling assuming an effective population size of roughly 2,500,000 (*e.g*. Duchen *et al*. 2013). Following the results of Uricchio & Hernandez (2014), we determined that under the RHH models of interest, a size reduction to 50,000 individuals should closely maintain the genetic variation of a non-rescaled population. Further, both algorithms provided in the cited paper yielded near identical scaled parameters when reducing the population size from 2,500,000 to 50,000 individuals. Because of this, we used the simpler algorithm 1 of Uricchio & Hernandez (2014). The main idea behind this rescaling method is that patterns of genetic diversity are maintained when population-scaled parameters *0 = 4N_e_μ., ρ = 4N_e_r*, and *γ = 2N_e_s* are fixed while *N_e_* is varied. As such, if we decrease *N_e_* =2,500,000 by 50X to *N_e_*=50,000, then *θ, ρ*, and *α* must be increased by 50X. The algorithm is laid out in step form below.

- Let *s*_0_ = α_0_/2*N*_0_*; r*_0_ *= ρ*_0_/4*N*_0_*; a = s*_0_/*L*_0_*r*_0_
- *γ*_1_*=γ*_0_
- *s*_1_ *= γ*_1_/2*N*_1_
- *r*_1_ *= s*_0_/a*L*_1_
- *λ*_1_ *= r*_1_*λ*_0_/*r*_0_

Here, the subscripts refer to before rescaling (subscript 0) and after rescaling (subscript 1). *s*_1_ is the selection strength, *N*_1_ is the population size, *r*_1_ is the per base pair per generation per chromosome recombination rate, and *L*_1_ is the simulated locus length.

We ran simulations under two different models of RHH, both of which were estimated from *D. melanogaster* data. Since the rate of adaptive substitutions and the average selective advantage are highly confounded in terms of their impact on diversity levels, we wanted to examine complementary models. The first model we chose to study is from Jensen *et al*. (2008). Here, the rate of incoming adaptive substitutions (*λ*) is low (*λ =* 4.2E-11; *2Nλ =* 2.1E-4) and the average strength of selection (*s* = 0.002; 2*Ns =* 10000) is high. The second model, from Andolfatto (2007), consisted of a high rate of adaptive substitutions (*λ =* 6.9E-10; 2*Nλ =* 3.45E-3) with a very low average selection strength (*s*=1.2E-5; 2*Ns =* 60). For both models, we used a mutation rate *μ* = 3.27E-9 (*4Nμ = 0.0327*) (Schrider *et al*. 2013) and a recombination rate *r* = 2.5E-8 (*4Nr =* 0.25). We also modeled gene conversion at a rate of 6.25E-8/bp/generation. The tract length of the gene conversion was drawn from a geometric distribution with a mean length of 518 base pairs (Comeron 2012). In forward simulations, it is not possible to directly specify the rate of adaptive substitutions. Instead, one must input the rate of beneficial mutations *v*. In the absence of interference among selected mutations, this can be derived using *λ* and the probability of fixation (Kimura 1962):

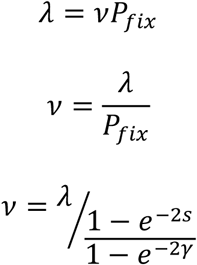

In our simulations, each beneficial mutation had its selection coefficient drawn randomly from an exponential distribution with a mean equal to about half of the rescaled selection strength. The means of these distributions were chosen such that the average selection strength of a fixed mutation is the rescaled selection strength, *s*_1_ (since the more strongly beneficial mutations drawn from this distribution are more likely to fix). The variation in selection coefficients helps to avoid the artificial scenario of interference between mutations with precisely identical fitness.

We wanted to analyze 10 kilobases (kb) from each simulation for demographic inference. Because a sweep can affect regions far from the target of a sweep, we simulated extra flanking regions for each side of the 10 kb that was used for the demographic inference, while analyzing only the middle region. For the common/weak sweep model, we simulated 480 base pairs on each side of the 10 kb for a total of 10,960 base pairs simulated. For the rare/strong model, we simulated 20 kb flanking loci for a total of 50 kb simulated. Using the formula (*2N_e_s*)*^−2r/s^* (Maynard-Smith and Haigh 1974, Barton 2010), we expect a 1% reduction in neutral diversity from a sweep 20 kb away under the rare/strong model, while under the common/weak model a sweep should reduce diversity 480 bp away by only 0.03%. Hence, these simulations should incorporate a large majority of sweep effects on neutral diversity predicted by the associated RHH models, while maintaining computational tractability. RHH simulation parameters are provided in Table 1.

**Table 1.**
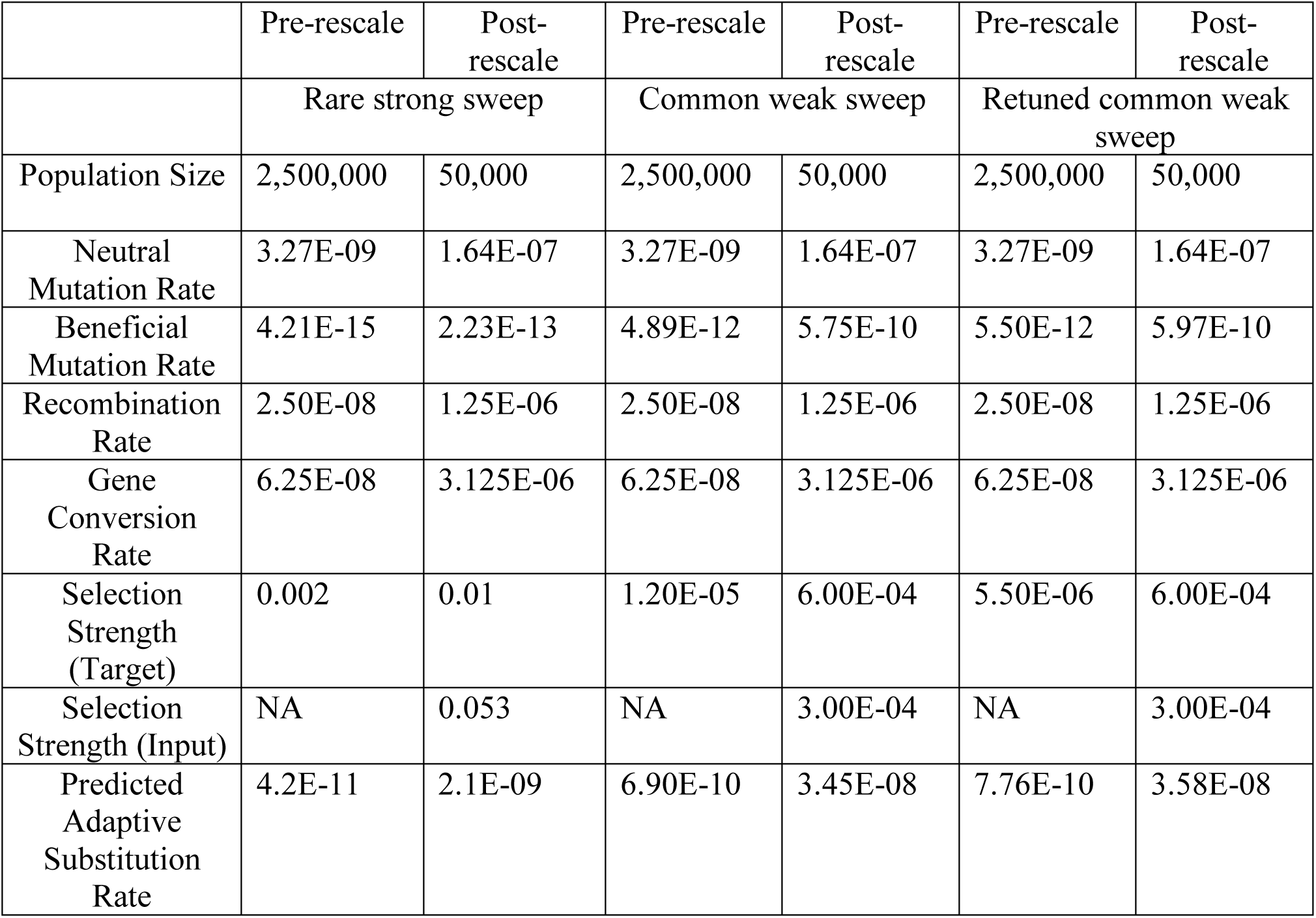
Summary of simulation parameters. The pre-rescale retuned common weak sweep parameters show the parameters for a full-size *Drosophila* population.

### Demographies simulated

Forward simulations require a burn-in period to generate appropriate genetic variation. Thus, both recurrent hitchhiking models were run for 500,000 (10*N_e_*) generations. These “trunk” simulations were then used for the demographic simulations. There are two relevant two-population demographic models that we were interested in. These demographies included a bottleneck model and an isolation with migration (IM) model. In our bottleneck model, the populations split and one population experiences a bottleneck. The parameters of the bottleneck model were taken from Thornton and Andolfatto (2006). In this model, a bottleneck occurs 0.0516 coalescent time units in the past (5,160 generations) and lasts for 0.042 coalescent time units (4,200 generations). During the bottleneck, the population decreases to 4.7% of its original population size. The IM model consisted of a population split *0.5N_e_* coalescent units in the past (25,000 generations) and subsequent migration of *2N_e_m* = 0.25. We simulated two different kinds of IM models, a “shared sweep” and a “private sweep” model. The “shared sweep” model allowed selective mutations to have an equal selective advantage if they migrated into the other population. The “private sweep” model multiplied s by −1 if it migrated into the other population, making the allele deleterious.

### Demographic Inference

To examine how recurrent hitchhiking affects demographic inference, we used δαδι version 1.6.3 (Gutenkunst *et al*. 2009) to estimate demographic model parameters. We first attempted to fit a two-epoch and three-epoch size change model to the trunk simulations (where there were no size changes simulated) to examine whether recurrent hitchhiking can misidentify demographic models. For the bottleneck simulations, we fit two and three parameter bottleneck models. The three parameter bottleneck model consisted of a population size reduction, a length of time as the reduced population size, and an instantaneous size change back to the original size that occurs some time in the past. In the two population bottleneck model, the length of the bottleneck is fixed and not optimized. We examined each bottleneck model both with and without fitting the ancestral size change models as well. In this way, we could better parse ancient parameters from more recent demographic parameters post population split. Finally, we tested the IM models with both shared and private selective sweeps. In these cases, the timing of the population split and the migration rate are estimated. As in the bottleneck cases, we tested both IM models with and without ancestral size changes. In total, 14 demographic models were investigated for both hitchhiking models and 11 demographic models were tested for the neutral simulations (since there is no shared/private sweep distinction in the neutral case).

We ran 1,000 simulations of both the common/weak RHH and the rare/strong RHH model. For any given demographic model tested, we randomly chose 50 simulations to generate a site frequency spectrum to run δαδι on. δαδι was then run 10 times on each SFS. The parameters of the δαδι run with the highest likelihood across the 10 runs were chosen as the inferred parameters. We repeated this process 200 times for each demography tested.

### Data Deposition

This study produced no empirical data. All scripts necessary to recapitulate the analyses presented can be found at http://github.com/jeremy-lange/RHH_project.

## Results

### Effects of Hill-Robertson Interference

In order to investigate the effects of recurrent hitchhiking on divergence and diversity in *Drosophila*, we performed forward simulations reflecting two published models of RHH representing relatively common/weak selection (Andolfatto 2007) and rare/strong selection (Jensen *et al*. 2008), respectively. Before proceeding with further analysis, we checked to see if our initial adaptive mutation rates based on these models were producing the prescribed rates of adaptive substitution, or if instead an important impact of interference must be accounted for. While the above studies assumed no interference between positively selected mutations, Andolfatto (2007) suggested that an interesting next step would be to examine how the presence of interference influences the observed versus expected adaptive substitution rate (*λ*) and selection coefficient of fixed beneficial substitutions (*s*) in the simulations.

Under a model of no interference, we can expect approximately *λLgP_fix_* adaptive substitutions as a product of the adaptive substitution rate at *L* sites across *g* generations and their probability of fixation. However, if multiple beneficial mutations are sweeping simultaneously, competition among sweeping haplotypes will lead fewer mutations to fix. Further, mutations that do fix will tend to have a higher selection strength than the input distribution of selective effects. The dynamics of the common/weak and rare/strong models that we tested are very different. In the rare/strong model, on average, we expect 554 beneficial mutations to occur per simulation (over the full 50 kb locus), with 52.2 fixing on average during the 500,000 generations. We expect one beneficial fixation approximately every 9,523 generations. It is unlikely, therefore, that any given beneficial mutation would experience interference from another. In the common/weak model, however, we expect 315,197 beneficial mutations per simulation. Under a model of no interference, we would expect on average 189 of these beneficial mutations to fix. This makes it very likely that more than one beneficial mutation would be sweeping at any given time. Thus, we expect interference to be more significant in the common/weak model relative to the rare/strong model.

To test this expectation, we ran both the common/weak and the rare/strong models as described in the methods section, using the published *λ* and *s* to generate input parameters. We tracked every mutation that fixed across all simulations and recorded the average selection coefficient. To accurately reflect the RHH models that we were simulating, our goal for our simulations was to approximately match the expected number of fixed beneficial mutations as described above. Across the 1000 simulations of the the rare/strong model, an average of 51.8 beneficial mutations fixed per simulation compared to an expectation of 52.2. For the common/weak RHH model, we found that, on average, 182 beneficial mutations fixed, corresponding to a modest 3.7% reduction in the expected number of adaptive fixations. We attribute this reduction to interference between positively selected mutations. In order to emulate the properties of the common/weak model, we increased the adaptive mutation rate by 3.7%. This recovered the desired rate of adaptive fixations, averaging 188 adaptive fixations per simulation.

We also conducted simulations to test whether our inclusion of gene conversion was crucial, and we found that it indeed had an important effect on the degree of Hill-Roberton interference. In simulations without gene conversion, the retuned common/weak model averaged 182 adaptive fixations. Thus, the absence of gene conversion reduced the rate of adaptive substitution by 3.3%.

We used the above retuned *v* parameter and ran 1,000 simulations as described in the methods section to more accurately reflect the common weak sweep RHH model. The output of these simulations were used for the simulation analyses detailed below. Note that for a non-rescaled *Drosophila* population, our adjusted RHH mutational parameters for the Andolfatto (2007) model would correspond to a beneficial mutation rate of *v* = 5.5E-12.

### Impacts of RHH on adaptive divergence

In these simulations, we can calculate the proportion of substitutions fixed by selection. In the rare/strong RHH model, 1.23% of all fixations are adaptive while in the retuned common/weak model, 17.5% of fixations are adaptive (Table 2). These proportions can be compared with the same quantity (*α*) estimated from extended McDonald-Kreitman analyses of empirical data (Smith and Eyre-Walker 2002). Our genome-wide analysis estimated that approximately 18.8% of substitutions in the *D. melanogaster* genome were driven to fixation by natural selection (Figure 1; Table S1). Our estimates of *α* are largely concordant with other studies (Andolfatto 2005, Begun *et al*. 2007, but see Mackay *et al*. 2012). Perhaps surprisingly, *α* estimates for two and three-fold synonymous site classes were very low and hence indicated evolutionary patterns similar to four-fold synonymous sites. It is unclear why these sites may not have been frequent targets of adaptive protein evolution and, further, it is unclear why *α* estimates did not converge to a particular range when the minimum allele frequency threshold was increased in the asymptotic McDonald-Kreitman method (Messer and Petrov 2013). For full results, see Table S1 for non-asymptotic *α* estimates with a simple 10% frequency threshold. Overall, this empirical analysis suggests that the common/weak RHH model seems more compatible with adaptive divergence estimates in *D. melanogaster*.

**Table 2.** The observed and expected proportion of substitutions driven by positive selection are shown. Expectations refer to theoretical predictions in the absence of interference.

| Model | Observed | Expected |
| --- | --- | --- |
| Rare/strong | 0.0123 | 0.0127 |
| Common/weak | 0.1702 | 0.1742 |
| Retuned common/weak | 0.1745 | 0.1796 |
| Empirical | 0.1884 |  |

**Figure 1:**
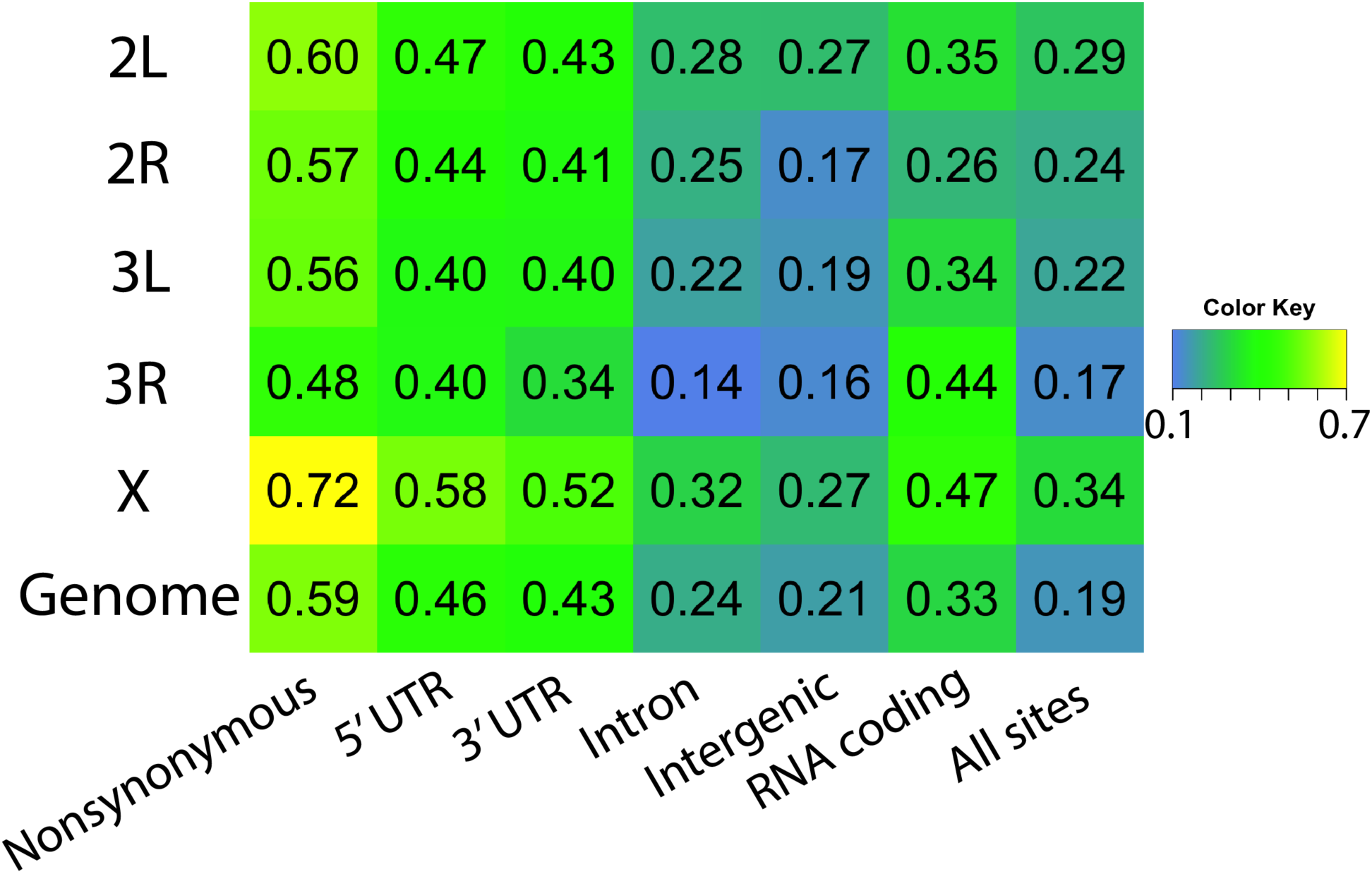
The estimated proportion of substitutions driven by positive selection (*α*), as estimated from *Drosophila* genomic data, is shown for each chromosome arm and site functional class, as well as the genome-wide average across all arms.

**Figure 2:**
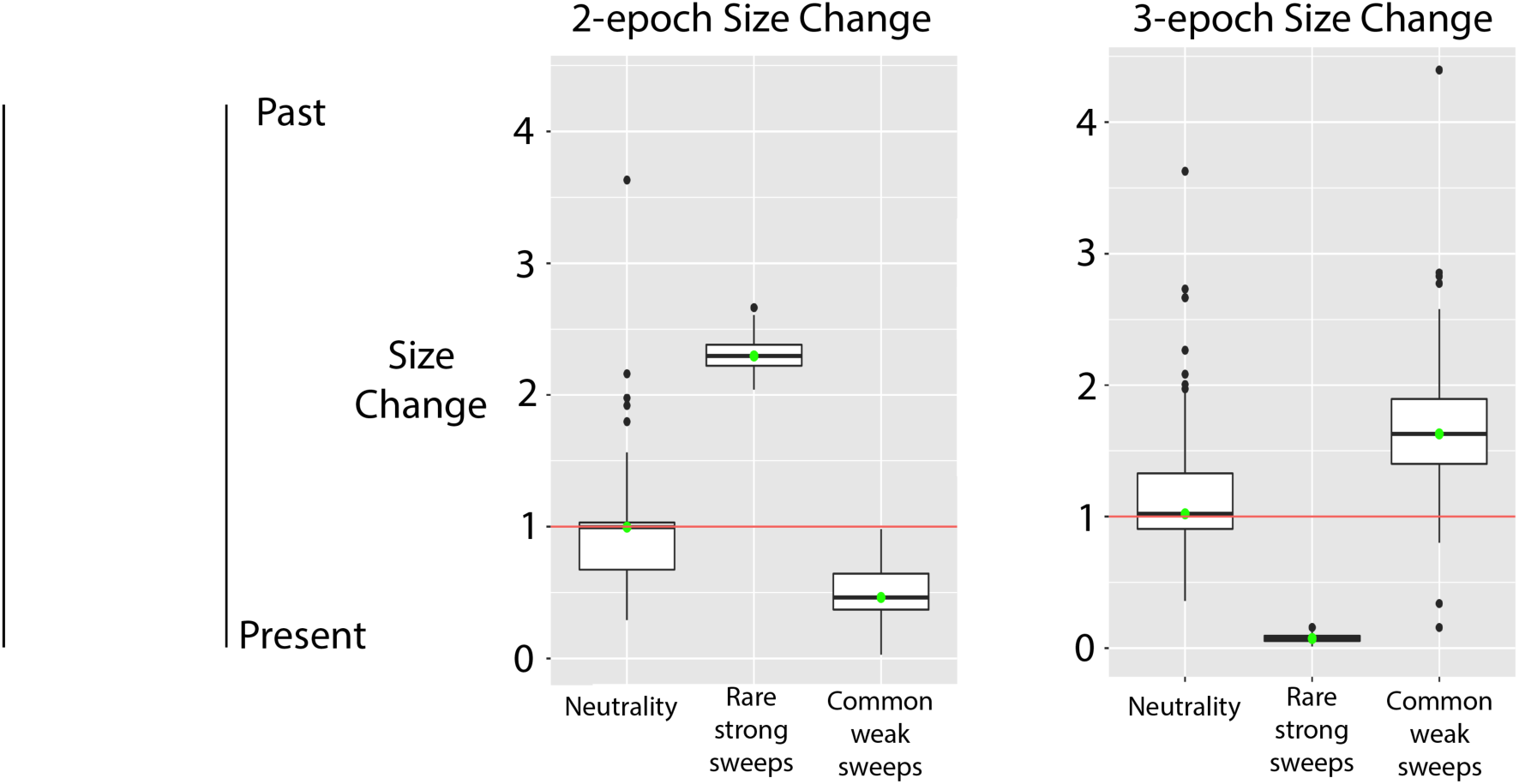
Falsely-inferred population size changes based on RHH simulations are illustrated. Population size change estimates are shown for the trunk simulations (no true size changes). In the two-epoch scenario, we allow δαδι to infer a single population size change. In the three-epoch scenario, we allow δαδι to infer a size change and require it to return to its original effective population size.

### Hitchhiking effects on demographic inference

The final goal of this study was to examine how models of recurrent hitchhiking affect inferences on demography. We tested whether selective sweep models involving substantial hitchhiking effects would violate assumptions of neutrality made by demographic inference tools and bias parameter estimation. We ran our simulations for 500,000 generations as a single population before we added a population split and distinct demographies. As previously demonstrated (Jensen *et al*. 2008), the rare/strong model entails a large impact on nucleotide diversity (*π* reduced by over 50% versus neutrality; Table S2). This model also generates a notable excess of rare alleles (Figure S1), in line with theoretical expectations (Braverman *et al*. 1995). By comparison, the common/weak model reduces *π* by just 15% and does not produce an excess of rare alleles.

For the “trunk” simulations, we first asked if δαδι would fit a model with population size changes over the true model of constant size. Here, we tested both two and three epoch size change models. The two epoch model consisted of two parameters to maximize: a single size change (reduction or expansion) that occurs some time in the past. The three epoch model has three parameters: a size change (reduction or expansion), a length of time at this new size, and some time in the past when the population recovers. The parameters of these models were most affected by the RHH models. Under neutrality, δαδι prefers a model of constant population size (the true model) for both the two and the three epoch models. In the two epoch size change model, δαδι infers a population expansion for the rare/strong sweep model, in line with the observed excess of rare alleles. For the three epoch model, a bottleneck ending *1.2N_e_* generations ago is instead favored. Qualititatively similar single-population results have recently been reported (Schrider *et al*. 2016). However, for common/weak sweep model, estimated population size changes are subtle (less than two-fold) and quite ancient (*>6N_e_* generations ago), in agreement with the lesser impact of RHH on genetic variation under this model.

Our primary interest was to assess whether the demographic parameters that relate recently-diverged populations are similarly biased by RHH. We therefore attempted to infer two distinct bottleneck models occurring after the 500,000 generation burn-in: a two and a three parameter model. Both models consisted of a population size contraction, a bottleneck length, and a length in time since recovery back to original population size. All three parameters are inferred in the three parameter model while bottleneck length is fixed and not inferred in the two parameter model. Results showed that in the two parameter bottleneck model, bottleneck strength was accurately inferred under neutrality and both selection models (Table S3). In the three parameter bottleneck case, bottleneck strength and bottleneck duration were accurately inferred under neutrality and in the common/weak RHH model, with little upward bias in the rare/strong RHH case (Figure 3; Table S3). Time since recovery was most affected by the presence of selection. Under neutrality, this parameter was accurately recapitulated for both the two and three parameter bottleneck models. Under selection, however, this parameter was overestimated by at least twofold. This result is in line with the fact that both post-bottleneck growth and selective sweeps leave behind an excess of rare alleles. Thus, if both are occurring in our simulation but δαδι assumes that only neutral events have occurred, it makes sense that this inference method is biased towards overestimating bottleneck recovery times in the presence of positive selection. Models were also run in which ancestral size change parameters and bottleneck parameters were estimated simultaneously, with qualitatively similar results (Table S4).

**Figure 3:**
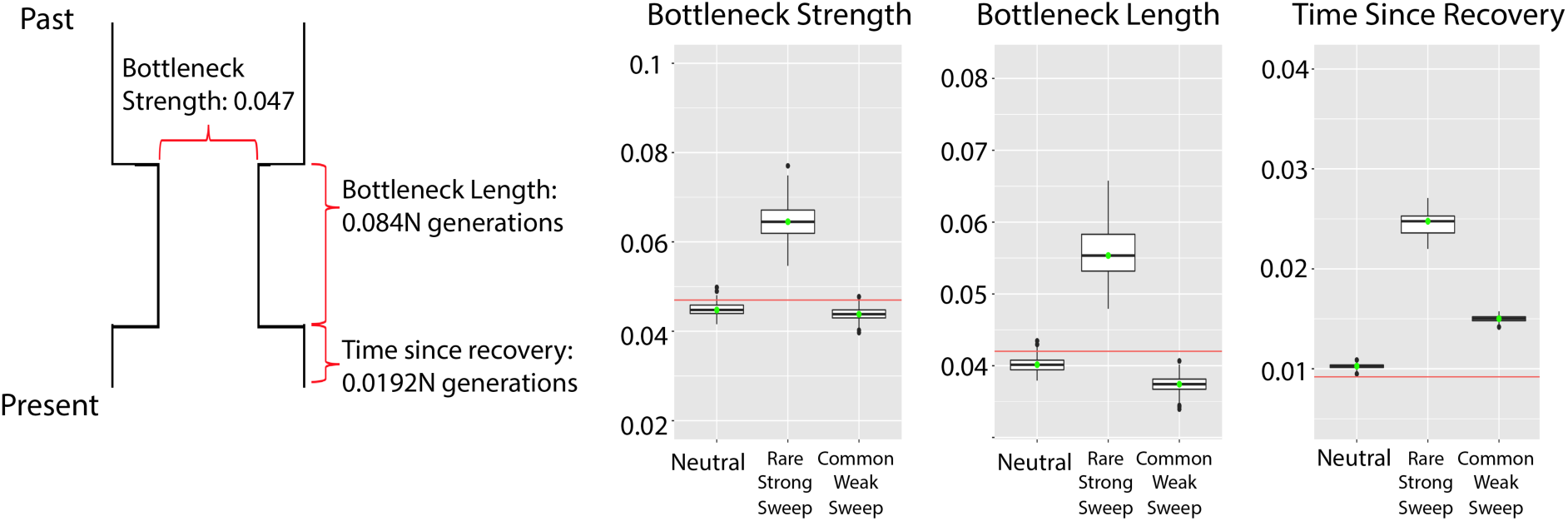
Demographic parameter estimates are shown for bottleneck simulations based on the true model shown on the left. Simulations with RHH showed relatively little bias for the duration and time since the bottleneck, but moderate bias for the time since recovery.

We simulated two distinct IM models: one with shared selective sweeps between populations, and one simulating local adaptation. For both cases, we estimated the time since the population split and the migration rate. It should be noted that the timescale simulated here, *0.5N_e_*, is much longer than the estimated divergence of any *D. melanogaster* populations (Duchen *et al*. 2013; Kern & Hey 2017). This scenario, therefore, should be viewed as an extreme scenario in terms of divergence time. Under neutrality, δαδι correctly estimated both the divergence time and the migration rate. Under both RHH models, however, these parameter estimates were biased in both the shared sweep and local adaptation cases. In the shared sweep model, δαδι overestimated migration rates, with less bias for divergence time (Figure 4; Table S3). In contrast, for the local adaptation case, the divergence time was greatly overestimated while the migration rate was less biased. As with the bottleneck models, we also made demographic inferences by combining the ancestral size change models with the recent divergence IM models, again with comparable results (Table S4).

**Figure 4:**
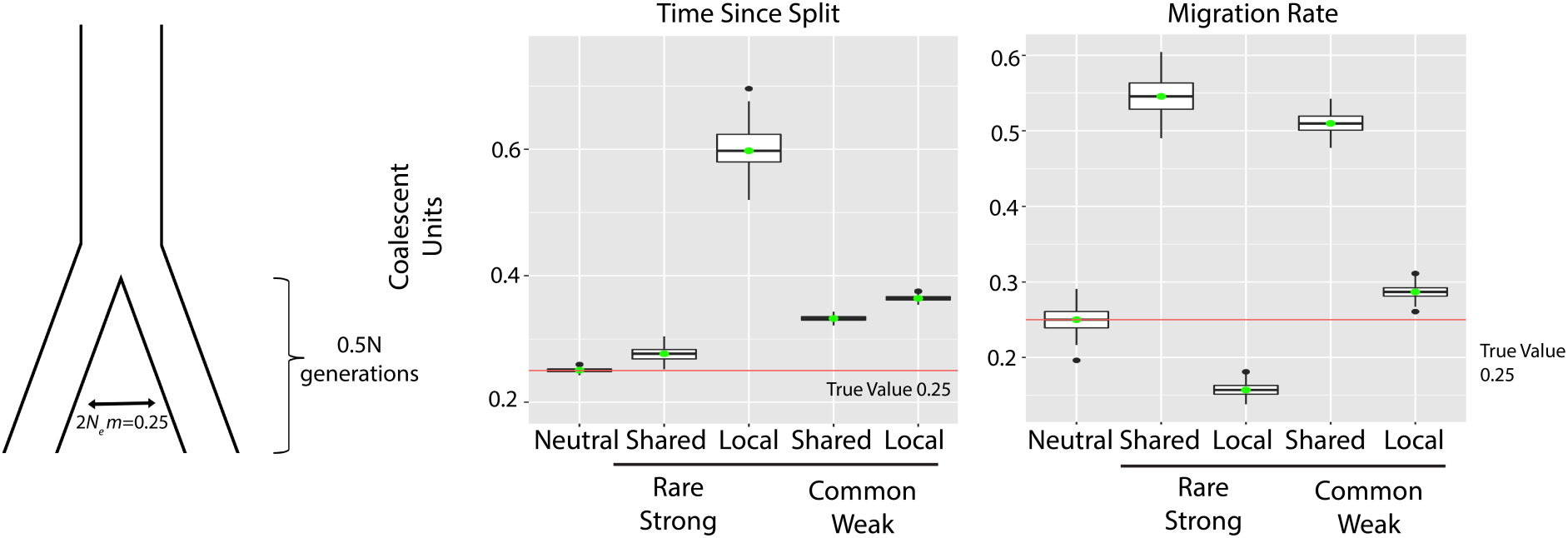
Demographic parameter estimates for simulations of the depicted isolation-migration model are shown. Some bias was observed for this relatively ancient population split time even under neutrality. However, differences from neutral estimates for RHH cases with shared sweeps or local sweeps were consistent with the effects of decreased or increased genetic differentiation, respectively.

## Discussion

Simulation provides us with flexible tools for studying the impact of selective sweeps on genetic diversity and divergence. In this study, we have examined two published RHH models and their consequences for divergence and demographic inference. We found that a model of common and weak positive selection appears to fit the genomic estimates of adaptive divergence better than a rare/strong sweep model, and that this favored model appears to entail a slight degree of Hill-Robertson interference. After retuning the common/weak model to yield adaptive substitution rates similar to the published parameters, we investigated the impact of both RHH models on demographic inference. Here, we confirmed that RHH can bias demographic parameters, but we found that the magnitude of such bias was greater for long term population parameters and lesser for more recent parameters.

Although we find agreement between the common/weak model and McDonald-Kreitman estimates of adaptive divergence, we do not argue for a specific quantitative RHH model in light of some important caveats. First, the model that gives *α* estimates in line with our genome-wide estimates is based on an analysis of nonsynonynous sites specifically (Andolfatto 2007). While this model may share traits in common with the true genomic RHH model for this species, it is best viewed as a qualitative example of the type of model compatible with this aspect of data. Second, we are assessing RHH models based on their general agreement with McDonald-Kreitman-based estimates of *α*, but such estimates can be biased depending on the history of population size change (Eyre-Walker 2002) and recombination rate change (Comeron 2014). Finally, estimates of *α* derived from RHH models depend not only on adaptive parameters, but also upon the *neutral* mutation rate that we use in simulations. The raw mutation rate may be somewhat higher than our simulated *μ* (Schrider *et al*. 2013; Huang *et al*. 2016). However, more than half of these mutations should be prevented from fixing by selective constraint (Halligan and Keightley 2006). If the *μ* that we use in our simulations is too high, the predictions for *α* may be too low, and vice versa. In spite of these quantitative uncertainties, we argue that our analyses provide general insight into the RHH models that are most plausible for this species.

Although this is not an inference study, our results suggest the importance of HRI in a common/weak RHH model like that of Andolfatto (2007). We found that this model led to a loss of approximately 3.7% of beneficial substitutions before retuning the adaptive mutation rate. These results raise the possibility that the effects of interference in *D. melanogaster* may be stronger than previously suggested (Castellano *et al*. 2016). In this 2016 study, the authors conclude that approximately 27% of beneficial substitutions are lost due to HRI. This estimate is based on the assumption that there is no interference in regions of recombination that exceed 2 cM/Mb. However, our results raise the prospect of slight interference in regions of high recombination (2.5 cM/Mb). Therefore, the genomewide effects of interference on the adaptive substitution rate may be somewhat stronger than previously estimated.

Our results also shed light on the impact of RHH on linked neutral variation and its consequences for demographic estimation. It is clear that strong effects of natural selection across the genome can violate the neutralist assumptions of typical demographic inference methods, whether due to background selection (Ewing and Jensen 2016) or selective sweeps (Schrider *et al*. 2016). Our results support the biasing effect of RHH on ancient parameter estimation and demographic model choice. This is in line with a previous study using single population simulations (Schrider *et al*. 2016). However, we show that there can be relatively less impact on recent parameters in two population models. Thus, such methods may retain utility even for populous species like *D. melanogaster*, but the biases that do exist should be borne in mind for this species and investigated for other taxa.

Our simulations emulate a population of *D. melanogaster*, a populous species with a compact genome. It has been shown that selection may be prevalent in large fly populations (Sella *et al*. 2009; Karasov *et al*. 2010; Langley *et al*. 2012). Further, due to the compactness of the genome, a selective sweep can affect relatively large regions. It is less clear how much of an impact natural selection would have on demographic inference in smaller *N_e_* species, such as humans. It is presumed that much of the genetic variation in the human genome is most affected by genetic drift instead of natural selection. Recent studies, however, have argued that an appreciable rate of adaptive substitution has shaped genetic variation in humans (Enard et al. 2014) and that soft sweeps play the dominant role in adaption in human evolution (Schrider and Kern 2017). Thus, even though theory suggests that demographic inference should be more accurate in a less populous species where the effects of natural selection should be lessened, further study is needed to delimit the parameter space in which RHH biases demographic inference.

The simulations in this study entail specific, important caveats. First, we simulated data reflecting the highly recombining portion of the *Drosophila* genome. The inclusion of low recombination regions would presumably exacerbate the effects of selective sweeps, HRI, and background selection on genetic variation and hence demographic estimates. Our simulations also did not model selective constraint. The biases we observed, therefore, would presumably be worse if analyzed sites had an excess of rare alleles due to deleterious variants. The RHH models that we simulated invoke an arbitrary distribution around a point estimate of *s*. Importantly, we do not know the true distribution of fitness effects in nature. Theoretical work has suggested that advantageous mutations may be exponentially distributed (Gillespie 1984, Orr 2003), as we have done in this study, although processes such as migration-selection balance may lead to departures from this prediction (Yeaman & Whitlock 2011). There are important caveats to such a distribution in regards to our inference. For instance, the size of the flanking region was determined by the point estimate of the selection strength. Any selective sweep with a selective strength substantially greater than the mean of the distribution will not fully be captured by our analyzed region.

The ABC method used in the Jensen study attempts to capture the locus-to-locus variance in genetic variation to infer adaptive mutation rate and selection strength. Such a method is highly dependent on the size of the genomic regions used in the study. Longer loci entail higher discriminatory power to distinguish between a rare/strong RHH model and a common/weak RHH model because common/weak selection reduces variance from one locus to the next even in small genomic windows. The loci used in the Jensen and Andolfatto studies were on average 680 base pairs, potentially biasing the ABC method of the Jensen study towards high values of *s*. The Andolfatto (2007) study utilized a McDonald–Kreitman approach to estimate *s*, which is not sensitive to the size of locus length. Our estimates of adaptive divergence are in accordance with the results of both the Andolfatto (2007) study and also a more recent study (Keightley **et al**, 2016). Using *D. melanogaster* polymorphism data, Keightley suggested a scaled selection strength of *N_e_s = 12*. This is very close to our common/weak model’s scaled selection strength of *N_e_s = 15* Further, the inferred probability that a new mutation is beneficial from the Keightley study (0.5%) is very close to the retuned estimate from this study (0.4%). Of course, while we draw these conclusions about weak/common RHH models, it is still possible that there could be a number of strong sweeps in the genome.

Our simulations only modeled complete sweeps from new mutations. It has been argued that this is not necessarily the dominant adaptive model in nature (Pritchard *et al*. 2010; Hernandez *et al*. 2011; Schrider and Kern 2017), so it is worth considering how other models of natural selection may alter conclusions drawn in this study. If there are two simultaneous soft sweeps, for instance, it is more likely that there exists a haplotype with both favorable variants prior to selection starting, reducing the impact of HRI. Likewise, because soft sweeps have more limited impacts on genetic variation (Pennings and Hermisson 2006), their impact on demographic estimation should be less severe as well.

It is also important to note that complete sweeps may only be a single component of Darwinian selection in nature. Other models of selection may impact genetic variation in fly populations with less effect on divergence, including fluctuating selection (Mustonen and Lässig 2010; Bergland *et al*. 2014) and diminishing selection (Vy *et al*. 2017). Thus, depending on the modes of positive selection that are prevalent in nature, the total impact of hitchhiking on genetic variation and demographic inference may be greater or lesser than simulated here – underscoring the need for further investigation of this topic.

## Acknowledgements

The UW-Madison Center for High Throughput Computing provided computational assistance and resources for this work. This work was funded by USDA Hatch grant WIS01900 to JDL.

## References

Andolfatto P. 2005. Adaptive evolution of non-coding DNA in *Drosophila*. Nature 437: 1149–52.

Andolfatto P. 2007. Hitchhiking effects of recurrent beneficial amino acid substitutions in the *Drosophila melanogaster* genome. Genome Res. 17: 1755–1762.

Begun DJ et al. 2007. Population genomics: whole-genome analysis of Polytene Chromosome Maps in *Drosophila* 1651 polymorphism and divergence in *Drosophila simulans*. PLoS Biol. 5: 2534–2559.

Bergland AO, Behrman EL, O’Brien KR, Schmidt PS, Petrov DA. 2014. Genomic evidence of rapid and stable adaptive oscillations over seasonal time scales in *Drosophila*. PLoS Genet. 10: e1004775.

Braverman JM, Hudson RR, Kaplan NL, Langley CH, Stephan W. 1995. The hitchhiking effect on the site frequency spectrum of DNA polymorphisms. Genetics 140: 783–796.

Castellano D, Coronado-Zamora M, Campos JL, Barbadilla A, Eyre-Walker A. 2016. Adaptive evolution is substantially impeded by Hill-Robertson interference in *Drosophila*. Mol. Biol. Evol. 33: 442–455.

Chevin LM, Hospital F. 2008. Selective sweep at a quantitative trait locus in the presence of background genetic variation. Genetics 180: 1645–1660.

Comeron JM, Ratnappan R, Bailin S. 2012. The many landscapes of recombination in Drosophila melanogaster. PLoS Genet. 8:e1002905.

Comeron JP. 2014. Background selection as baseline for nucleotide variation across the *Drosophila* genome. PLoS Genet. 10: e1004434.

Duchen P, Zivkovic D, Hutter S, Stephan W, Laurent S. 2013. Demographic inference reveals African and European admixture in the North American *Drosophila melanogaster* population. Genetics 193: 291–301.

Enard D, Messer PW, Petrov DA. 2014. Genome-wide signals of positive selection in human evolution. Genome Res. 24: 885–895.

Ewing GB, Jensen JD. 2016. The consequences of not accounting for background selection in demographic inference. Mol. Ecol. 25: 135–141.

Eyre-Walker A. 2002. Changing effective population size and the McDonald-Kreitman test. Genetics 162: 2017–2024.

Gerrish P, Lenski R. 1998. The fate of competing beneficial mutations in an asexual population. Genetica 102–103: 127–144.

Gillespie JH. 1984. Molecular evolution over the mutational landscape. Evolution 38: 1116–1129.

Gutenkunst RN, Hernandez RD, Williamson SH, Bustamante CD. 2009. Inferring the joint demographic history of multiple populations from multidimensional SNP frequency data. PLoS Genet. 5: e1000695.

Haller BC, Messer PW. 2016. Slim 2: Flexible, interactive forward genetic simulations. Mol. Biol. Evol. 34: 230–240.

Haller BC, Messer PW. 2017. asymptoticMK: A web-based tool for the asymptotic McDonald-Kreitman test. G3: Genes, Genomes, Genetics. 7(5): 1569–1575.

Halligan DL, Keightley PD. 2006. Ubiquitous selective constraints in the *Drosophila* genome revealed by a genome-wide interspecies comparison. Genome Res. 16: 875–884.

Hernandez RD et al. 2011. Classic selective sweeps were rare in recent human evolution. Science 331: 920–924.

Hill WG, Robertson A. 1966. The effect of linkage on limits to artificial selection. Genet. Res. 8: 269.

Huang W et al. 2016. Spontaneous mutations and the origin and maintenance of quantitative genetic variation. eLife 5:e14625.

Jensen J D, Thornton KR, Andolfatto P. 2008. An approximate Bayesian estimator suggests strong, recurrent selective sweeps in *Drosophila*. PLoS Genet. 4: e1000198.

Kaplan NL, Hudson R, Langley C. 1989. The “hitchhiking effect” revisited. Genetics 123: 887–899.

Karasov T, Messer SW, Petrov DA. 2010. Evidence that adaptation in *Drosophila* is not limited by mutation at single sites. PLoS Genet. 6: e1000924.

Kern AD, Hey J. 2017. Exact calculation of the joint allele frequency spectrum for isolation with migration models. Genetics 207: 241–253.

Kimura M. 1962. On the probability of fixation of mutant genes in a population. Genetics 47(6): 713–719.

Lack JB et al. 2015. The *Drosophila* genome nexus: A population genomic resource of 623 *Drosophila melanogaster* Genomes, including 197 from a single ancestral range population. Genetics 199: 1229–1241.

Langley CH et al. 2012. Genomic variation in natural populations of *Drosophila melanogaster*. Genetics 192: 533–598.

Mackay T F et al. 2012. The *Drosophila melanogaster* genetic reference panel. Nature 482: 173–178.

Maynard Smith J, Haigh J. 1974. The hitch-hiking effect of a favourable gene. Genet Res. 23: 23–35.

McDonald JH, Kreitman M. 1991. Adaptive protein evolution at the Adh locus in *Drosophila*. Nature 351: 652–654.

Messer PW, Petrov DA. 2013. Frequent adaptation and the McDonaldKreitman test. Proc. Natl. Acad. Sci. USA 110: 8615–8620.

Mustonen V, Lässig M. 2010. Fitness flux and ubiquity of adaptive evolution. Proc. Natl. Acad. Sci. USA 107: 4248–4253.

Orr HA (2003) The distribution of fitness effects among beneficial mutations. Genetics, 163, 1519–1526.

Pennings PS, Hermisson J. 2006. Soft sweeps III: the signature of positive selection from recurrent mutation. PLoS Genet. 2: e186.

Pool JE et al. 2012. Population genomics of Sub-Saharan *Drosophila melanogaster:* African diversity and non-African admixture. PLoS Genet. 8: e1003080.

Pritchard JK, Pickrell JK, Coop G. 2010. The genetics of human adaptation: hard sweeps, soft sweeps, and polygenic adaptation. Curr. Biol. 20: R208–R215.

Schrider DR, Houle D, Lynch M, Hahn MW. 2013. Rates and genomic consequences of spontaneous mutational events in *Drosophila melanogaster*. Genetics 194: 937–954.

Schrider DR, Shanku AG, Kern AD. 2016. Effects of linked selective sweeps on demographic inference and model selection. Genetics 204: 1207–1223.

Schrider DR, Kern AD. 2017. Soft Sweeps are the dominant mode of adaptation in the human genome. Mol. Biol. Evol. 34: 1863–1877.

Sella G, Petrov DA, Przeworski M, Andolfatto P. 2009. Pervasive natural selection in the *Drosophila* genome? PLoS Genet. 5: e1000495.

Smith NGC, Eyre-Walker A. 2002. Adaptive protein evolution in *Drosophila*. Nature 415: 1022–1024.

Stanley CE, Kulathinal RJ. 2016. Genomic signatures of domestication on neurogenetic genes in *Drosophila melanogaster*. BMC Evol. Biol. 16: 6.

Stephan W, Wiehe TH, Lenz MW. 1992. The effect of strongly selected substitutions on neutral polymorphism: analytical results based on diffusion theory. Theor. Popul. Biol. 41: 237–254.

Thornton K, Andolfatto P. 2006. Approximate Bayesian inference reveals evidence for a recent, severe bottleneck in a Netherlands population of *Drosophila melanogaster*. Genetics 172(3): 1607–19.

Uricchio LH, Hernandez RD. 2014. Robust forward simulations of recurrent hitchhiking. Genetics 197(1):221–36.

Vy HMT, Won YJ, Kim Y. 2017. Multiple modes of positive selection shaping the patterns of incomplete selective sweeps over African populations of *Drosophila melanogaster*. Mol. Biol. Evol. 34: 2792–2807.

Wiehe T, Stephan W. 1993. Analysis of a genetic hitchhiking model, and its application to DNA polymorphism data from *Drosophila melanogaster*. Mol. Biol. Evol. 10: 842–854.

Yeaman S, Whitlock MC. 2011. The genetic architecture of adaptation under migration-selection balance. Evolution 65: 1897–1911.

